# Growth adaptation of *gnd* and *sdhCB Escherichia coli* deletion strains diverges from a similar initial perturbation of the transcriptome

**DOI:** 10.1101/342683

**Authors:** Douglas McCloskey, Sibei Xu, Troy E. Sandberg, Elizabeth Brunk, Ying Hefner, Richard Szubin, Lydia Beeken, Adam M. Feist, Bernhard O. Palsson

**Affiliations:** Department of Bioengineering, University of California - San Diego, La Jolla, CA 92093, USA.; Novo Nordisk Foundation Center for Biosustainability, Technical University of Denmark, 2800 Lyngby, Denmark.

**Keywords:** adaptive laboratory evolution, *gnd*, *sdhA*, *sdhB*, *sdhC*, *sdhD* gene knockouts, casual mutations, multi-omics analysis, multi-omics integration, systems biology, *E. coli*

## Abstract

Adaptive laboratory evolution (ALE) has emerged as a new approach with which to pursue fundamental biological inquiries and, in particular, new insights into the systemic function of a gene product. Two *E. coli* knockout strains were constructed: one that blocked the Pentose Phosphate Pathway (gnd KO) and one that decoupled the TCA cycle from electron transport (*sdhCDAB* KO). Despite major perturbations in central metabolism, minimal growth rate changes were found in the two knockout strains. More surprisingly, many similarities were found in their initial transcriptomic states that could be traced to similarly perturbed metabolites despite the differences in the network location of the gene perturbations and concomitant rerouting of pathway fluxes around these perturbations. However, following ALE, distinct metabolomic and transcriptomic states were realized. These included divergent flux and gene expression profiles in the *gnd* and *sdhCDAB* KOs to overcome imbalances in NADPH production and nitrogen/sulfur assimilation, respectively, that were not obvious limitations of growth in the unevolved knockouts. Therefore, this work demonstrates that ALE provides a productive approach to reveal novel insights of gene function at a systems level that cannot be found by observing the fresh knockout alone.

## Introduction

Biological systems contain many overlapping metabolic and regulatory networks that contribute to robustness against perturbations. Systems robustness to perturbation is due to regulatory adjustments that can coordinate re-routing of flux by adjusting enzyme levels to compensate for metabolic pathway disruption ^1–4^. A systems level understanding of how regulatory adjustments are made can be gained by knocking out a gene product and studying the downstream metabolic and regulatory shifts that occur. Genes encode two major metabolic reactions were removed from a pre-evolved strain of *E. coli* K-12 MG1655. The two reactions were GND (*gnd*, 6-phosphogluconate dehydrogenase) and SUCDi (genes *sdhA*, *sdhB*, *sdhC*, and *sdhD* corresponding to the enzyme Succinate Dehydrogenase). GND carries out the final NADPH generating step of the oxidative Pentose Phosphate Pathway (oxPPP) to produce D-ribulose-5-phosphate (ru5p-D), which is a necessary precursor for nucleotide biosynthesis. SUCDi converts succinate (succ) to fumarate (fum) in the TCA cycle and also directly charges and donates quinones to the electron transport chain (ETC) via Complex II, thus directly coupling the TCA cycle to respiration.

Only minimal changes in growth rate were found when disabling the GND and SUCDi reactions. It has been shown previously that disrupting the *gnd* gene resulted in non appreciable changes to growth rate, but induces major changes in fluxes through the PPP and TCA cycle ^1–3^. Similar observations have also been made of *zwf* mutants ^4,5^, which demonstrate that metabolic networks can rapidly adjust enzyme level and flux without an appreciable cost to growth rate. The same ability to re-route fluxes has not been demonstrated for disruptions to the TCA cycle as has been done for the PPP. Importantly, the question of whether or not the immediate regulatory and flux shifts were the most optimal has not been explored.

The adaptive response to gene loss can be studied using adaptive laboratory evolution (ALE). ALE is an experimental method that introduces a selection pressure (e.g., growth rate selection) in a controlled environmental setting ^6–8^. Using ALE, organisms can be perturbed from their evolutionary optimized homeostatic states, and their re-adjustments can be studied during the course of adaptation to reveal novel and non intuitive gene product functions and interactions ^9^. When applied to KO strains in *E. coli*, it has been shown that the growth rate loss from the removal of key metabolic genes can be overcome through the selection of beneficial mutations ^10–12^. The rate of accumulation of compensatory adaptation is often associated with the amount of initial growth rate lost ^13^. The relationship between rate of compensatory adaptation and relative growth rate change would imply that little to no compensatory adaptation would be expected to occur when the environmental or genetic change resulted in minimal growth rate loss. The relationship between rate of compensatory adaptation and relative growth rate change would also imply that little to no changes in the regulatory or metabolic network would be found post-evolution.

In this study, the evolutionary drivers in the absence of significant growth rate loss were explored. Starting from a pre-optimized *E. coli* K-12 MG1655 strain, two major metabolic perturbations were inflicted that blocked the oxPPP and decoupled the TCA cycle from the ETC. Minimal changes in growth rate were found in the KO strains and evolved KO endpoints. Detailed omics analysis demonstrated that despite minimal changes in growth rate, massive changes in metabolic flux, gene expression, and metabolite levels occurred in both KO and evolved strains. Interestingly, many of the changes found in the regulatory network were shared by both knockout strains despite major differences in the location of the metabolic perturbation. Investigation of mutations in the evolved end points indicated that in the absence of a significant change in growth rate, selection pressures existed that resulted in major adaptations in regulatory and metabolic networks. These mutations led to divergent adjustments at the regulatory and metabolic networks in both knockout strains. These results demonstrated even when genetic perturbations induce little cost to growth rate, major optimizations to regulatory and metabolic network structure can be found after evolution.

## Results

### Strain selection: starting with a growth optimized strain

In order to eliminate any confounding effects between the adaptation to the growth conditions used in the experiment and the loss of a gene product, we chose a wild-type *E. coli* K-12 MG1655 strain previously evolved under glucose minimal media at 37°C ^14^ (denoted as “Ref”). Based on this rationale, this reference strain has been used as the basis for several ALE studies ^15, 16^.

### Blockage of the oxidative PPP and decoupling of the TCA cycle from the ETC resulted in minimal fitness loss

GND (*gnd*, 6-phosphogluconate dehydrogenase) and SUCDi (genes *sdhA*, *sdhB*, *sdhC*, and *sdhD* corresponding to the enzyme Succinate Dehydrogenase) were removed from Ref to generate strain uGnd and uSdhCB (denoted “unevolved *gnd* and *sdhCB* knockout strain”). The initial growth rate of uGnd and uSdhCB were minimally changed (9* and 6* decrease in growth rate, respectively) (Fig. 1D). 3 uGnd and 3 uSdhCB stains from independently inoculated starting cultures were simultaneously evolved on glucose minimal media at 37°C in an automated ALE platform ^14, 16^ denoted “evolved *gnd* and *sdhCB* knockout strains” or “eGnd and eSdhCB”. A non-significant and minimal increase in final growth rate was found in all endpoints of the eSdhCB and eGnd lineages (ave±stdev 3±4, 5±3* increase in growth rate) compared to the unevolved knockout strains (uSdhCB and uGnd lineages), respectively (Student’s t-test, pvalue<0.05).

**Fig. 1.**
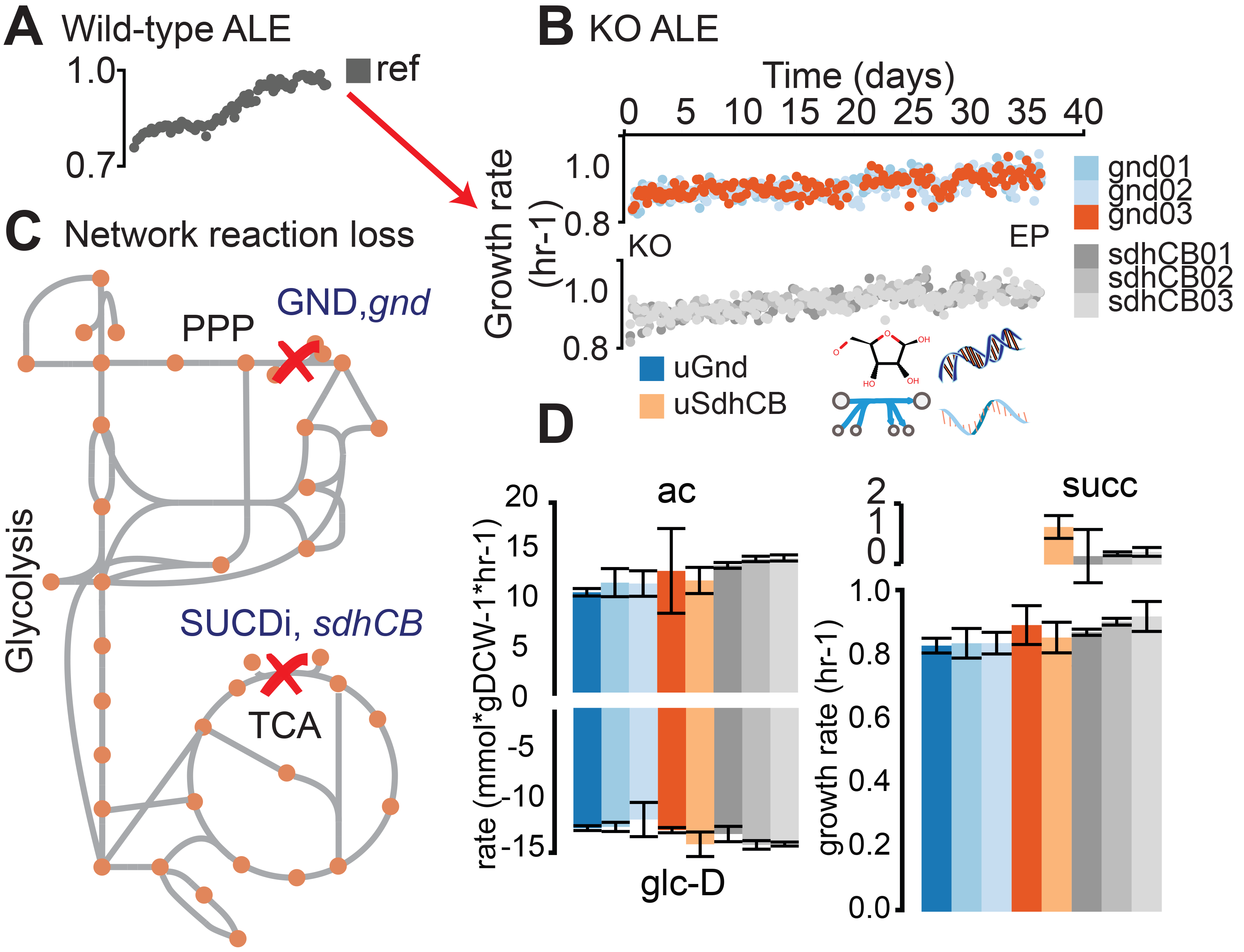
Evolution of knockout (KO) strains from a pre-evolved (i.e., optimized) wild-type strain. A) Wild-type (wt) *E. coli* (MG1655 K-12) was previously evolved on glucose minimal media at 37°C^14^. An isolate from the endpoint of the evolutionary experiment was selected as the starting strain for subsequent KOs of *gnd* and *sdhCB* and adaptive laboratory evolution (ALE). B) Adaptive laboratory evolution trajectories of the evolved knockout lineages. -Omics data collected included metabolomics, fluxomics, physiology, DNA resequencing, and transcriptomics. C) 6-phosphogluconate dehydrogenase (GND) and Succinate Dehydrogenase (SUCDi) were disabled by the gene KOs. GND converts 6-phosphogluconate (6pgc) to ribulose 5 phosphate (ru5p-D) in the Pentose Phosphate Pathway (PPP), and SUCDi converts succinate (succ) to fumarate (fum) in the TCA cycle. D) Growth rate and uptake and secretion rates for unevolved KO (uGnd and uSdhCB) and evolved KOs (eGnd and eSdhCB).

Genome-scale models were used to compute the flux map in Ref. The results indicated that GND and SUCDi represent two of the highest flux reactions when Ref was grown on glucose minimal media (data not shown). The minimal changes in growth rate after the loss of these two reactions were surprising given that massive flux rerouting would have to take place in each of the uKO strains. Detailed-omics analysis of all uGnd, eGnd, uSdhCB, and eSdhCB strains were carried out to better understand the drivers for flux rerouting in the absence of substantial growth rate loss.

### Blocked flux through the oxidative PPP in uGND overflowed into the ED pathway

6-phosphogluconate dehydrogenase (GND) encoded by *gnd* catalyzes the decarboxylation of D-gluconate-6-phosphate (6pgc) to D-ribulose-5-phosphate (ru5p-D) while regenerating NADPH. Loss of *gnd* created a flux bottleneck at the 6pgc node, which resulted in a massive buildup of 6pgc and depletion of ribose-5-phosphate (r5p) in uGnd. 6pgc was 3.3 log2 fold higher and r5p was −0.6 log2 fold lower in concentration in uGnd than Ref. The relatively large proportion of flux compared to Ref remained through the first steps of the OxPPP, but the flux that would flow through GND instead spilled over into the Entner-Doudoroff (ED) pathway in uGnd (Fig. 2A, Table S6). The spillover amounted to a 1.3 log2 fold higher absolute flux in ED in uGnd compared to Ref.

**Fig. 2.**
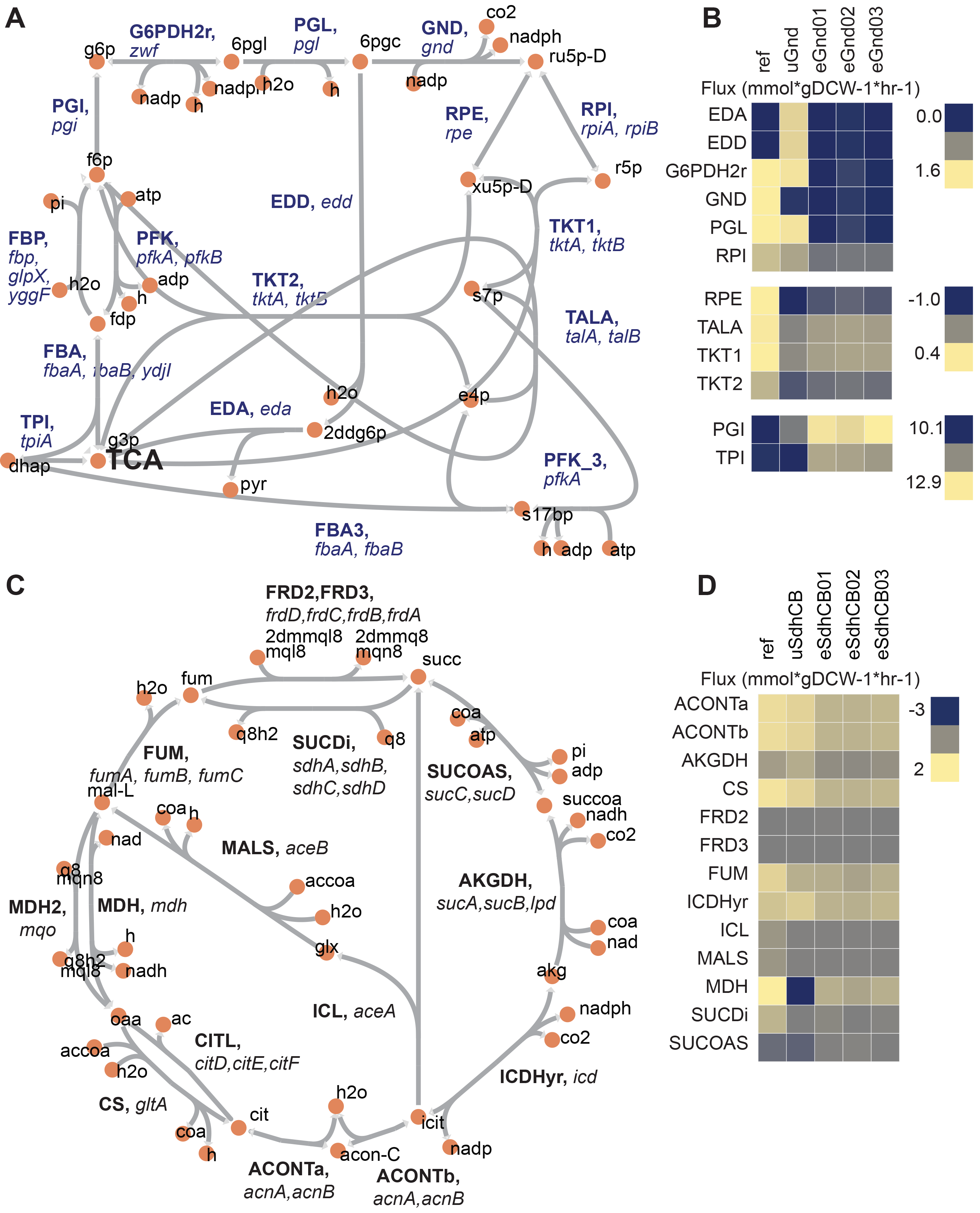
Flux maps of *gnd* and *sdhCB* knockout strains at the vicinity of the network perturbation. A) Network diagram of upper glycolysis, the oxidative pentose phosphate pathway (oxPPP), and the non oxidative pentose phosphate pathway (nonOxPPP). B) Measured fluxes for Ref, uGnd, and eGnd strains. C) Network diagram of the TCA cycle. D) Measured fluxes for Ref, uSdhCB, and eSdhCB.

An overall decrease in OxPPP flux was also accompanied by an increase in upper glycolytic flux by necessity of balancing the glucose-6-phosphate (g6p) node (Fig. 2A, Table S6). Thus, a 1.1 log2 fold higher absolute flux through upper glycolysis was found in uGnd compared to Ref. The increase in glycolytic flux also increased the levels of glycolytic metabolites (i.e., glucose-6-phosphate, fructose-6-phosphate, fructose 1,6-bisphosphate, and dihydroxyacetone phosphate, Table S3). For example g6p was 1.3 log2 fold higher in uGnd compared to Ref.

The r5p node was regenerated by re-routing flux through the non oxidative Pentose Phosphate Pathway (NonOxPPP) (Fig. 2A, Table S6). These flux changes resulted in a flip in flux direction through the transketolase enzymes (TKT1 and TKT2) as well as the transaldolase enzyme (TALA).

### Purine and pyrimidine biosynthetic pathways were downregulated due to transcription factor activation in both uGnd and uSdhCB strains

The building blocks for purines and pyrimidines is r5p, generated from the PPP. Activity of the majority of purine and pyrimidine biosynthetic genes are feedback inhibited by end products including AMP, GMP, and UMP ^17^. In addition, gene expression of the majority of purine and pyrimidine biosynthetic genes are negatively regulated by the PurR transcription factor (TF), which is itself activated by hypoxanthine (hxan) ^18, 19^.

Interestingly, all *de novo* purine and pyrimidine biosynthetic genes were downregulated in uGnd and uSdhCB and subsequently restored to levels similar to Ref in eGnd and eSdhCB strains (Table S4). Down regulation of purine and pyrimidine biosynthetic genes appears non-intuitive because the levels of the purine and pyrimidine building block, r5p, and flux through the PPP were at dramatically different levels in uGnd and uSdhCB strains. However, the levels of the PurR TF activator, hxan, was significantly elevated in both uGnd and uSdhCB strains. hxan was 4.2 and 4.6 log2 fold higher in uGnd and uSdhCB, respectively, compared to Ref. Given the down regulation of *de novo* purine and pyrimidine biosynthetic genes and non-significant change in *purR* expression (Table S4), the similar expression profiles of purine and pyrimidine genes in uGnd and eGnd appeared to be driven by similar levels of the PurR TF activator, hxan. hxan was restored to levels similar to Ref in eGnd and eSdhCB strains.

### Decoupling of the TCA from the ETC in uSdhCB resulted in flux cycling and increased flux towards nitrogen assimilation

SUCDi catalyzes the conversion of succinate (succ) to fumarate (fum) while contributing electrons directly to the ETC. Removal of *sdhA, sdhB, sdhC*, and *sdhD* resulted in a massive build-up of succ that was largely excreted into the medium (Fig. 1D). Even with the loss of Succinate Dehydrogenase (SUCDi), a small amount of conversion from succ to fumarate (fum) was found (Fig 2C,D, Table S6). 0.09 mmol*gDCW-1*hr-1 absolute flux (or −1.4 log2 fold change in uSdhCB compared to Ref) was found through SUCDi. A massive increase in gene expression of the *frdABCD* operon, which encodes the Fumarate Reductase (FRD) enzyme, was found ^20^. FRD is an iron-sulfur protein that is optimally active under anaerobic conditions ^20^. However, even when oxygen is present, FRD is able to catalyze the oxidation of succinate at a ratio of 1:1.5 compared to reduction of fumarate ^21^. For reference, SUCDi catalyzes the oxidation of succinate at a ratio of 25:1 compared to reduction of fumarate ^21^. The large expression levels of FRD and the massively elevated levels of succ indicated that the small amount of conversion from succ to fum was either due to mass action or suboptimal activity of FRD. Succ was 6.3 log2 fold higher in uSdhCB compared to Ref.

A larger, yet depleted, amount of flux exiting the fum node through FUM appeared to be driven primarily by recycling of fum from peripheral metabolism (Fig. 2C,D, Table S6). Flux from succ to fum accounted for less than a quarter of the flux through FUM. In addition, a reduced amount of flux through the glyoxylate shunt was found (−0.7 log2 fold less in uSdhCB compared to Ref).

Surprisingly, an overall increase in flux entering the TCA cycle was found in uSdhCB (Fig. 2C,D, Table S6). The increase in flux was driven by an increase in flux through Phosphoenolpyruvate Carboxylase (PPC) and maintenance of flux through Citrate Synthase (CS) leading to a 0.3 log2 fold change in Isocitrate Dehydrogenase (ICDHyr) flux in uSdhCB compared to Ref. Approximately half of the flux leaving the alpha-ketoglutarate (akg) node was directed towards nitrogen metabolism.

A cycle between flux entering the TCA through PPC and flux leaving the TCA through the Malic Enzymes (MEs) was also found (Fig. 2C,D, Table S6). The cycle was made possible by a complete reversal of Malate Dehydrogenase (MDH) from 1.3 mmol*gDCW-1*hr-1 to -1.4 mmol*gDCW-1*hr in uSdhCB. This cycle appeared to deplete levels of oxaloacetate (oaa) based on the measured levels of L-aspartate (asp-L) while maintaining the levels of Malate (mal-L) (Table S3).

### Common expression profiles in uGnd and uSdhCB in TCA cycle genes were driven by transcription factor and two component system activation by metabolites

Despite loss of enzymatic activity and major flux re-routing in distal network locations, expression profiles of TCA cycle genes were found to be remarkably similar (Fig. 3). In particular, the expression levels of genes that encode fumarate reductase (*frdA*, *frdB*, *frdC*, and *frdD*) and succinate dehydrogenase (*sdhA*, *sdhB*, *sdhC*, and *sdhD*) were almost identical in uGnd and uSdhCB strains. In fact, expression of the *sdh* operon in uGnd was downregulated to near the levels measured in the KO uSdhCB (Fig. 3, Table S4).

**Fig. 3.**
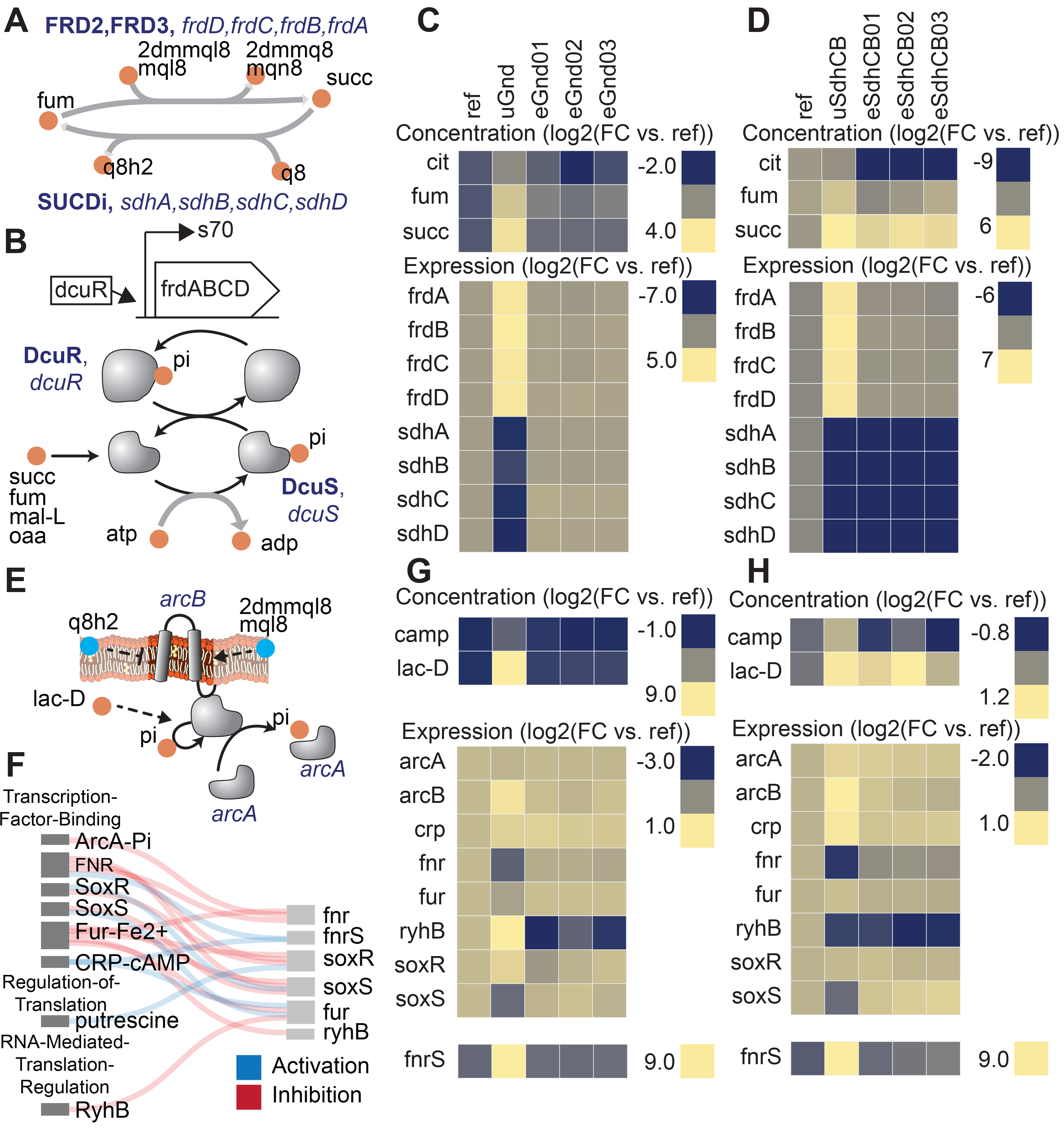
Perturbations in separate network locations yielded similar expression states in the uKOs due to similar metabolite levels. Removal of the succinate dehydrogenase complex (sdhCB), which decoupled the TCA cycle from oxidative phosphorylation, and removal of 6-phosphogluconate dehydrogenase (gnd), which re-routed flux through upper glycolysis, the ED pathway, and the pentose phosphate pathway (PPP), resulted in similar expression profiles of TCA cycle genes as a result of increased levels of intracellular four-carbon acids (Panels A-D). A) Reactions catalyzed by succinate dehydrogenase (SUCDi) and fumarate reductase (FRD2, FRD3) in the TCA cycle. B) Schematic of the *frd* operon and regulation by DcuR. Also shown is a schematic of the dcuRS two component system^22^. Elevation in four-carbon acids (e.g., succinate, fumarate, malate, and oxaloacetate) were detected by the dcuRS two-component system in the uKO strains. Phosphorylated DcuR activated expression of the fumarate reductase operon genes. Metabolite levels of fumarate (fum), succinate (succ), and citrate (cit), and expression levels of fumarate reductase (*frdA*, *frdB*, *frdC*, *frdD*) and succinate dehydrogenase (*sdhA*, *sdhB*, *sdhC*, *sdhD*) genes for gnd (Panel C) and sdhCB (Panel D). The similar metabolite levels in the uGnd and uSdhCB activated a network response that resulted in the upregulation of *frdABCD* genes in uGnd and uSdhCB, and downregulation of *sdhABCD* genes in uGnd. De-coupling of the TCA cycle from oxidative phosphorylation triggered an attenuated anaerobic response in gnd and sdhCB that involved a complex interaction of TFs ArcA, CRP, SoxR, SoxS, Fur, and Fnr, and small RNAs *fnrS* and *ryhB* (Panels E-H). E) Regulatory schematic of the signal transduction cascade triggered by the oxidized status of the membrane bound quinones ubiquinone (q8 and q8h2) and menaquinols (mql8, mqn8, 2dmmql8, and 2dmmq8), and anaerobic metabolites (e.g., lac-D). F) Regulatory interaction diagram between the different regulators. Metabolite and expression profiles of key components involved in the regulatory cascade for gnd (Panel G) and sdhCB (Panel H). Note the similar downregulation of *fnr* in response to ArcA activation through increased levels of lac-L and changes in the oxidized status of the membrane bound quinones, the upregulation of *fnrS* in response to activation of CRP-cAMP through increased levels of cAMP, and the downregulation of *soxS* in uGnd and uSdhCB.

Upregulation of expression of the *frd* operon in uGnd and uSdhCB was found to be attributed to a common elevation in four-carbon acids (i.e., succinate, fumarate, and malate) (Fig. 3B-D). Expression of the *frd* operon was most likely activated by the DcuR TF, which was most likely phosphorylated and activated by two component (TC) system pair DcuS in response to elevations in four-carbon acids ^22^. succ, fum, and mal-L were 3.7, 2.9, and 1.21 log2 fold higher in uGnd compared to Ref, respectively, and 6.3, 0.7, and −0.1 log2 fold change in uSdhCB, respectively, compared to Ref. Downregulation of the *sdh* operon in uGnd and uSdhCB was found to be attributed to an attenuated anaerobic response that involved a complex interaction of TFs ArcA, CRP, SoxR, SoxS, Fur, and Fnr, and small RNAs *fnrS* and *ryhB* (Fig. 3E-H). A shift in the oxidized status of the membrane bound quinones ubiquinone (q8 and q8h2) and menaquinones (mql8, mqn8, 2dmmql8, and 2dmmq8), and anaerobic metabolite (i.e., lac-D) most likely triggered the ArcAB TC, which phosphorylated and activated the ArcA TF. The ensuing regulatory cascade following ArcA activation culminated in the similar downregulation of *fnr.* Coupled with a common activation of CRP-cAMP through increased levels of cAMP resulted in the downregulation of *soxS* and upregulation of *fnrS.* Unique to uGnd was the activation of *rhyB*, the small regulatory RNA, which may have attributed to the downregulation of *sdhABCD* genes.

### Differences in uGnd and uSdhCB OxPPP and TCA cycle fluxes and derived metabolites were found

In contrast to uGgnd, an overall increase in flux through the OxPPP was found in uSdhCB. In contrast to uSdhCB, an increase in flux through the lower half of the TCA cycle was found in uGnd (and also in eGnd strains) (Table S6). In particular, the increased flux through ICDHyr was most likely to compensate for the loss of NADPH generation in the oxPPP. NADPH was - 1.3 log2 fold lower in uGnd compared to Ref. However, flux at the akg node was not significantly diverted towards nitrogen assimilation as was the case in uSdhCB. The insignificant divergence of flux towards nitrogen assimilation is consistent with previous work that has found an upregulation in TCA cycle flux towards succ in response to disruption of the *gnd* gene ^3, 23^. These differences, among several others, set the stage for physiological advantages to be had through evolution and compensatory mutations.

### Mutations at the *pntA* promoter boosted NADPH levels in eGnd

Intergenic mutations that increased gene expression of the NADP(H) binding component of the insoluble pyridine nucleotide transhydrogenase (THD2pp) were found in eGnd strains (Fig. 4). Specifically, single nucleotide polymorphisms (SNP) mutations in the vicinity of the *pntA* transcription start site (TSS) were found in all eGnd strains. THD2pp is composed of two subunits encoded by *pntA* and *pntB.* The former contains the NADP(H) binding domain while the latter contains the proton pumping transmembrane domain ^24^. Significantly elevated expression of *pntA* was found in all eGnd strains. The expression levels of *pntA* were 2.5, 2.1, and 3.2 log2 fold higher in eGnd01, 02, and 03, respectively, compared to uGnd. THD2pp has been shown to provide an important source of NADPH in *E. coli* ^25^, and it is highly likely that increased expression of *pntA* contributed to restoring the levels of NADPH in the eGnd strains.

**Fig. 4.**
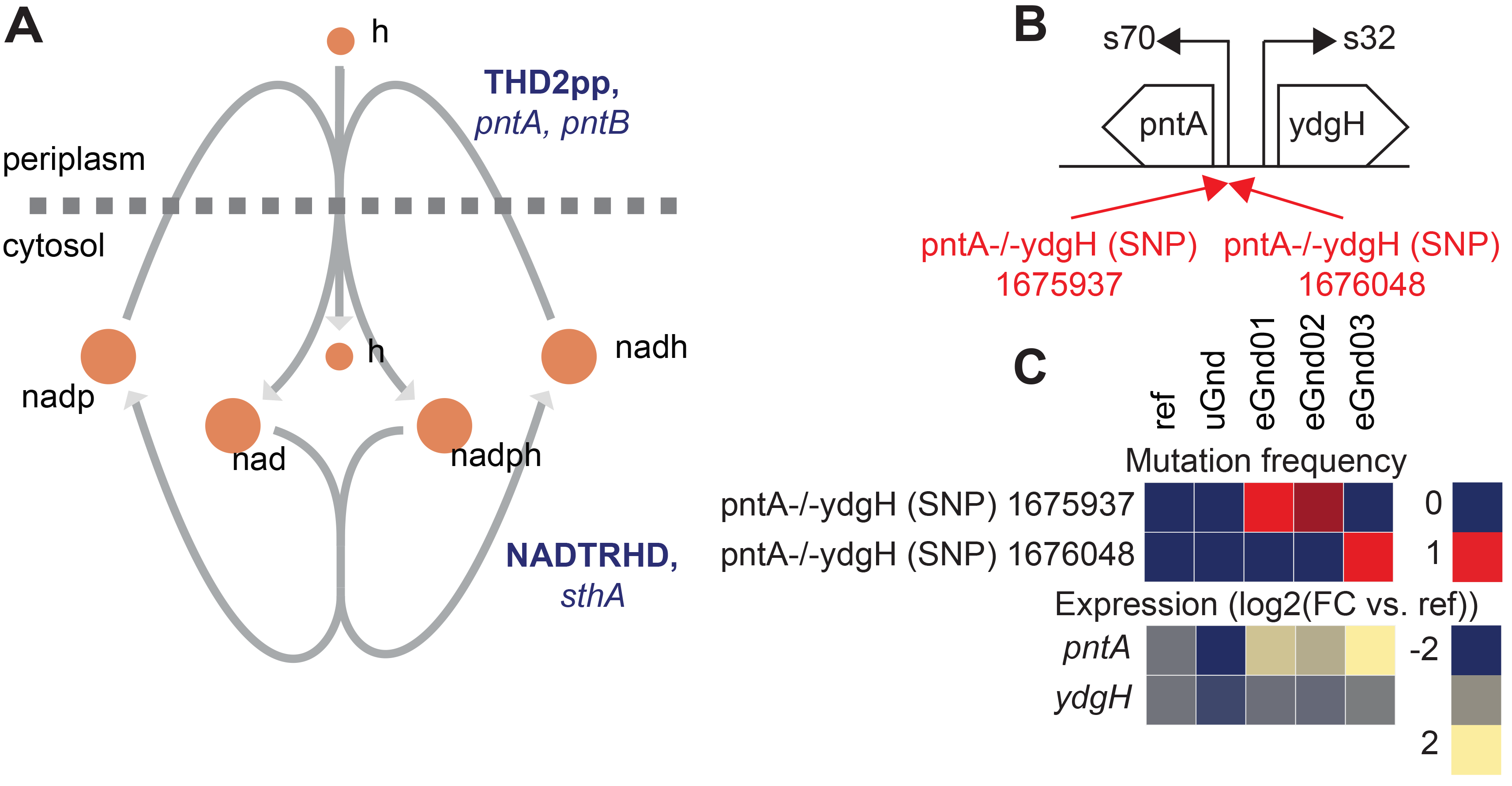
Intergenic mutations that increased gene expression of the NADP(H) binding component of the insoluble pyridine nucleotide transhydrogenase in eGnd strains. A) network diagram of the insoluble (THD2pp) and soluble (NADTRHD) pyridine nucleotide transhydrogenase. THD2pp has been shown to provide an important source of NADPH in *E. coli* ^25^. The insoluble pryidine nucleotide transhydrogenase is composed of two subunits encoded by *pntA* and *pntB.* The former contains the NADP(H) binding domain while the latter contains the proton pumping transmembrane domain ^24^. B) Operon diagram of *pntA.* Mutations that affected gene expression of *pntA* were found in all eGnd strains. The 1675937 SNP is 62 nucleotides downstream of the *pntA* transcription start site (TSS) and the 1676048 SNP is 49 nucleotides downstream of the pntA TSS. Mutations are annotated in red. C) Mutation frequency and expression levels of *pntA* and *ydgH.* Note that all eGnd strains significantly elevated expression of *pntA.*

### Mutations affecting nitrogen assimilation in eSdhCB strains may have coordinated increased flux out of the TCA cycle with nitrogen regulation

The loss of SUCDi in uSdhCB created a bottleneck in the TCA cycle which led to an elevated amount of flux exiting the TCA in the steps leading to the production of succinate. As described above, this would lead to an increased amount of flux directed towards nitrogen assimilation, which resulted in a significant elevation of key nitrogen sensing metabolites, akg and gln-L (p-value<0.05). akg and gln-L were 1.2 and 1.0 fold higher in uSdhCB compared to Ref. Several mutations were found in eSdhCB strains that altered enzymatic activity of metabolic genes and key regulators in nitrogen assimilation pathways that potentially balanced increased flux with nitrogen generation. These included mutations in *allD* and *glnD* that are described below.

A mutation in the active site of ureidoglycolate dehydrogenase (URDGLYCD) in eSdhCB01 was found that potentially provided an auxiliary means to metabolize excess glyoxylate (glx) and/or balance nitrogen levels (Fig. 5). The *allDCE* operon encodes genes involved in converting allantoin (alln) to glyoxylate (glx) and oxaluric acid (oxur) ^26^ (Fig. 5B). oxur can then be broken down to oxymate (oxam), and carbamoyl phosphate (cbp); the latter provides a source of ATP and nitrogen ^26^ (Fig. 5B). The *allDCE* operon is positively regulated by AllS, whose gene expression is negatively regulated by AllR ^27^ (Fig. 5A). *allA* and *allB* are also negatively regulated by AllR ^27^ (Fig. 5A). Glyoxylate enzymatically inhibits AllR repression of *allS, allA*, and *allB ^21^* (Fig. 5A). A deletion (DEL) mutation was found in eSdhCB01 in the active site of URDGLYCD that would most likely affect binding of NAD and other substrates (Fig. 5C). Interestingly, the gene expression profiles of all allonoin related genes are consistent with high glyoxylate levels in uSdhCB, which were found to be 3.3 log2 fold higher in uSdhCB compared to Ref (Fig. 5D, Table S3, Table S4). Given the high amounts of glyoxylate and also increased flux towards nitrogen metabolism, the mutation in eSdhCB01 may have conferred a fitness advantage by providing an additional means to metabolize glyoxylate and/or balance the levels of ammonia and nitrogen in the cell.

**Fig. 5.**
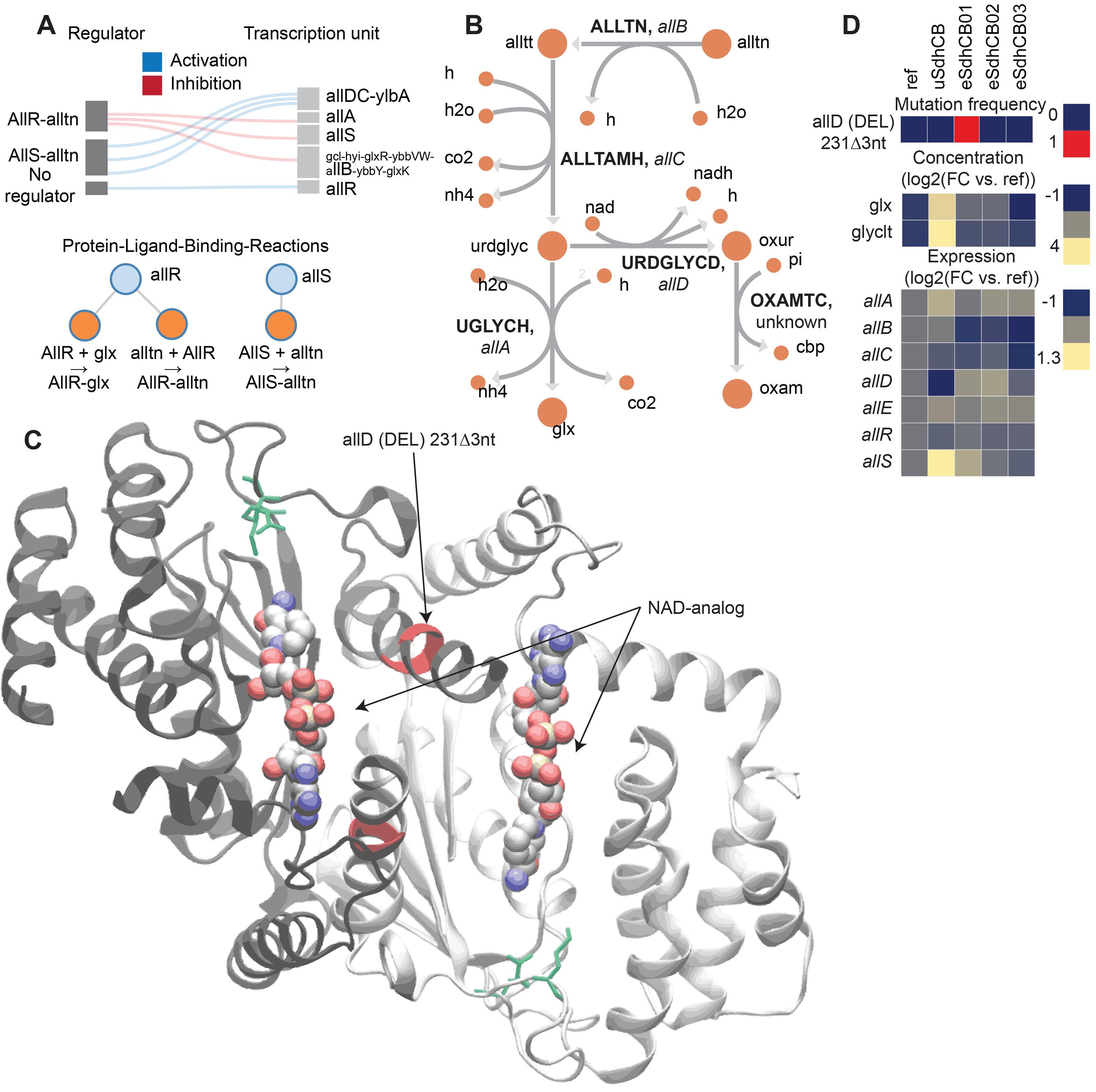
A mutation in the active site of ureidoglycolate dehydrogenase (URDGLYCD) in eSdhCB01 that potentially provided an auxiliary means to metabolize excess glyoxylate (glx) and/or balance nitrogen levels. A) Regulatory diagram and protein-ligand-binding reactions. The *allDCE* operon is positively regulated by AllS, whose gene expression is negatively regulated by AllR ^27^. *allA* and *allB* are also negatively regulated by AllR ^27^. Glyoxylate enzymatically inhibits AllR repression of *allS, allA*, and *allB ^27^.* B) Network diagram of allantoin assimilation. C) Crystal structure of AllD [Reference]. Mutations are highlighted in red. Active site residues are highlighted in green. An NAD-analog where NAD would bind is shown. Note that the deletion (DEL) mutation in eSdhCB01 would affect active site residues, and most likely binding of NAD and other substrates. D) Mutation frequency, metabolite levels, and gene expression of components involved in allantoin assimilation. Note that gene expression profiles are consistent with high glyoxylate levels in uSdhCB.

Mutations were also found in *glnD*, which encodes the primary nitrogen status sensor PII uridylyl-/deuridylyl-transferase, that fixed in eSdhCB02 and that overtook a majority of the population in eSdhCB03 (Fig. 6). Nitrogen assimilation is heavily regulated in *E. coli* (see ^28^ for a review) (Fig. 6A-B). PII senses nitrogen levels primarily through the concentration of L-glutamine (gln-L). Increased levels of gln-L increase the deuridylylation activity of glnD and decrease the uridylyltransferase activity of glnD ^29^. PII-ump stimulates deadenylation of GLNS via glnE while PII stimulates adenylation of GLNS ^28, 30^; the removal of amp enhances GLNS activity ^28, 30^. The mutations in *glnD* were located in the ACT 1 and 2 domains (Fig. 6C), which are believed to play a direct role in binding and sensing gln-L ^29, 31^. Given the increased flux towards nitrogen assimilation out of the TCA cycle and elevated gln-L levels (Table S3, Table S6), the mutations potentially provided a fitness advantage by altering regulation of nitrogen assimilation in eSdhCB strains.

**Fig. 6.**
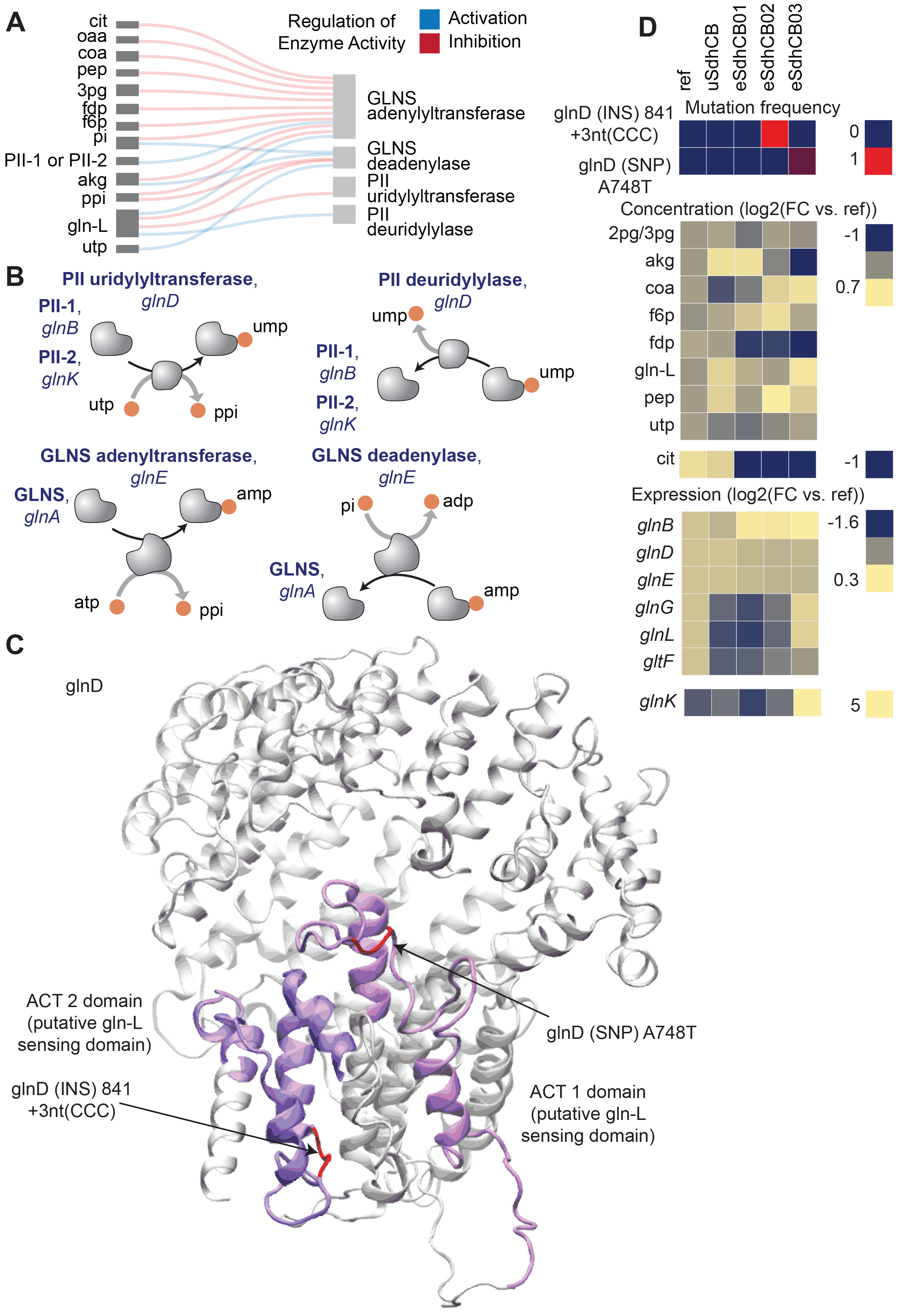
Mutations that potentially affect regulation of nitrogen assimilation in eSdhCB strains. Mutations in *glnD*, which encodes the primary nitrogen status sensor PII uridylyl-/deuridylyl-transferase, fixed in eSdhCB02 and overtook a majority of the population in eSdhCB03. A) Regulatory diagram of allosteric regulation of PII uridylyl-/deuridylyl-transferase and GLNS adenylyltransferase/GLNS deadenylase. B) Protein-protein interaction diagram of PII uridylyl-/deuridylyl-transferase and GLNS adenylyltransferase/GLNS deadenylase. Increased levels of gln-L increase the deuridylylation activity of glnD and decrease the uridylyltransferase activity of glnD ^29^. PII-ump stimulates deadenylation of GLNS via glnE while PII stimulates adenylation of GLNS ^28, 30^; the removal of amp enhances GLNS activity ^28, 30^. C) Crystal structure of PII uridylyl- /deuridylyl-transferase [Reference]. Mutations are highlighted in red. The ACT 1 and 2 domains are highlighted in dark and light magenta, respectively. The ACT 1 and 2 domains are believed to play a role in sensing the intracellular levels of gln-L ^29, 31^. Note that the mutations were located in the ACT 1 and 2 domains. D) Mutation frequency, metabolite levels, and gene expression levels of components involved in *E. coli* nitrogen assimilation.

### Altered TCA cycle flux perturbed sulfur metabolism gene expression in uSdhCB

Major perturbations in sulfur metabolic pathway gene expression were found in uSdhCB (Fig. S2). The sulfur metabolic pathway converts sulfate (so4), asp-L, L-serine (ser-L) and Succinyl-CoA (succoa) to L-cysteine (cys-L), which is then converted to L-methionine (met-L). The pathway is also regulated enzymatically and transcriptionally by multiple intermediates and end-products of the pathway. Major changes included a significant decrease in gene expression of *cysDNC* and *cysJIH* operons in uSdhCB (Fig. 2A-B, E-F). *cysDNC* and *cysJIH* operons encode enzymes involved in sulfate reduction, and are controlled by the TF CysB. A significant decrease in gene expression of *thrA* and *metL* were found (Fig S2C-D). *thrA* and *metL* encode the fused aspartate kinase/homoserine dehydrogenase 1 and 2 (ASPK and HSDy), respectively. ASPK and HSDy catalyze the conversion of asp-L to L-aspartyl-4-phosphate (4pasp) and L-aspartate-semialdehyde (aspsa) to L-homoserine (hom-L) ^32, 33^. *metL* gene expression is repressed by high met-L levels ^34^. The levels of met-L were 0.82 log2 fold higher in uSdhCB compared to Ref.

Gene expression changes in the eSdhCB strains resulted in an increased flux towards hom-L biosynthesis (Fig. S2D). ASPK, ASAD, and HSDy flux increased by 0.7, 0.7, and 0.9 log2 fold in eSdhCB strains, respectively, compared to Ref. A significant decrease in gene expression of *cysE, cysK*, and *cysM* genes were found (Fig. S2E-F). *cysE, cysK*, and *cysM* encode enzymes involved in L-cysteine biosynthesis. In conjunction with *cysE, cysK*, and *cysM* down regulation, a significant decrease in gene expression of *metB, metC, metH*, and *metE* was also found. *metB, metC, metH*, and *metE* encode enzymes involved in met-L biosynthesis.

In contrast to the down regulation of the majority of sulfur metabolic genes, a significant increase in gene expression of *metA* was found (Fig. S2E-F). *metA* encodes the first step in *de novo* L-methionine biosynthesis catalyzed by homoserine O-succinyltransferase (HSST). One of the substrates of HSST, succinyl-CoA (succoa), is derived from the TCA cycle and two steps removed from SUCDi. Interestingly, other inputs into sulfur metabolism, asp-L and ser-L, were perturbed due to changed metabolic flux (Table S3). These results indicated that the perturbed gene expression of the majority of sulfur metabolic genes could be attributed to changed pathway flux and levels of key signalling metabolites.

### Intergenic mutations were found that enhanced expression of sulfur metabolic genes in eSdhCB strains

Expression of sulfur metabolic pathway genes were restored to levels similar to Ref in all eSdhCB strains. The restoration of expression was most likely due to restoration of key regulator metabolites including met-L (Table S3). However, significantly elevated expression of the *cysDNC* operon in eSdhCB03 was found that most likely resulted from an intergenic mutation (Fig. 7). Two intergenic mutations were found that targeted the RNAP binding site (eSdhCB02) and the CysB binding site (eSdhCB03) (Fig. 7A). *cysDNC* is positively regulated by the CysB TF. *cysD* and *cysN* encode subunits of sulfate adenylyltransferase (SADT2) which converts sulfate (so4) to adenosine 5’-phosphosulfate (aps) (Fig. 7B). *cysC* encodes adenylyl-sulfate kinase (ADSK) which converts aps to 3’-phosphoadenylyl-sulfate (paps) (Fig. 7B). The eSdhCB03 was found to have significantly higher expression levels of *cysDNC* than all other strains (Fig. 7C).

**Fig. 7.**
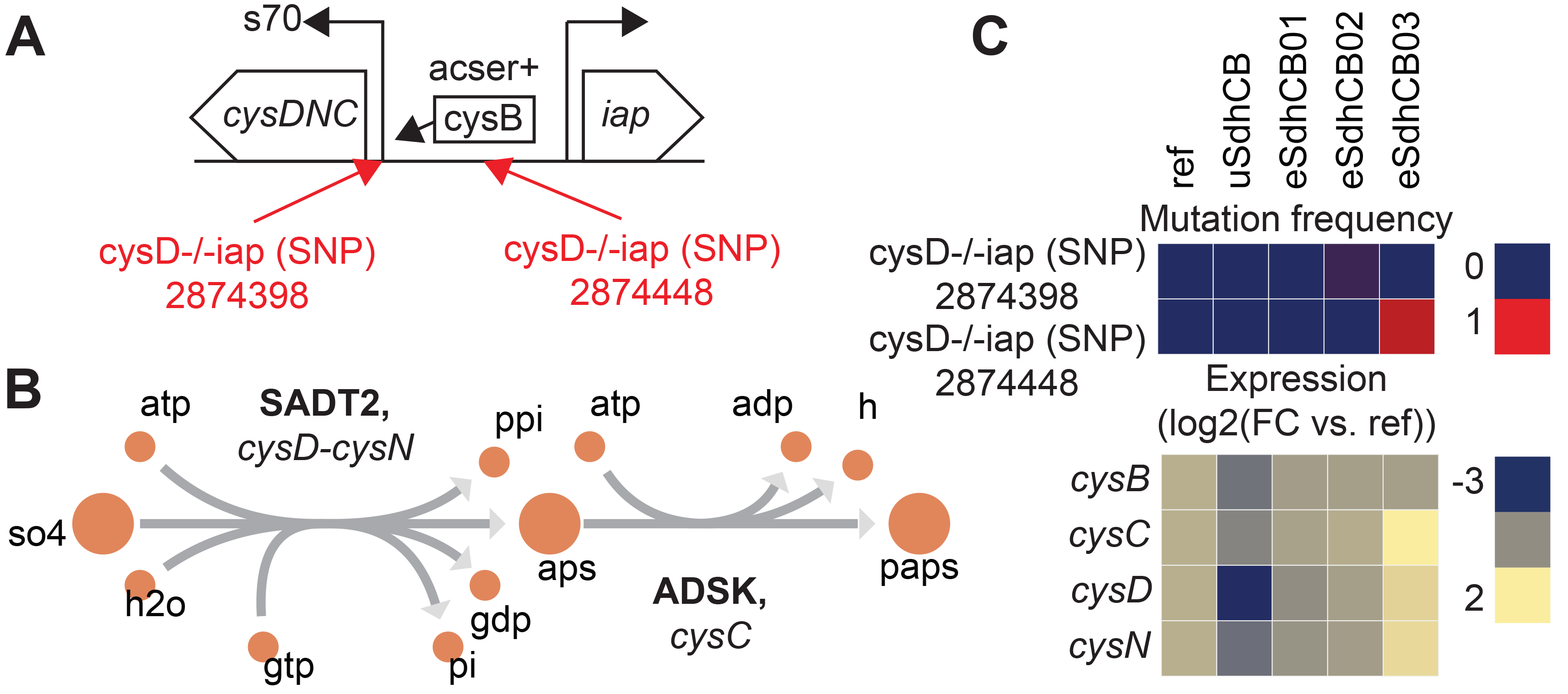
Intergenic mutations that enhance expression of the *cysDNC* operon in eSdhCB strains. A) Operon diagram of the *cysDNC* and *iap* operons. *cysDNC* is positively regulated by the transcription factor CysB, and encodes enzymes involved in sulfate reduction. Mutations are annotated in Red. The 2874398 SNP is located at the RNAP binding site. The 2874448 SNP is located at the CysB binding site. B) Network schematic of the reactions encoded by *cysD, cysN*, and cysC. *cysD* and *cysN* encode subunits of sulfate adenylyltransferase (SADT2) which converts sulfate (so4) to adenosine 5’-phosphosulfate (aps). *cysC* encodes adenylyl-sulfate kinase (ADSK) which converts aps to 3’-phosphoadenylyl-sulfate (paps). C) Mutation frequency and expression levels for genes involved in expression of the *cysDNC* operon. Note that eSdhCB03 has significantly higher expression levels of *cysDNC* than all other strains.

### Mutations in AKGDH decreased TCA cycle flux and reduced succinate secretion in all eSdhCB strains

Succinate secretion was significantly reduced in almost all eSdhCB strains compared to uSdhCB (Fig. 1D). Overall flux from citrate (cit) to succ was also significantly reduced in all eSdhCB strains (Fig. 2D). A major contributor to reduced TCA cycle flux and succinate secretion were mutations that affected 2-oxoglutarate Dehydrogenase (AKGDH) expression and enzymatic activity leading to significantly depressed flux in all eSdhCB strains (Fig. 8). AKGDH is a multimer, composed of multiple subunits encoded by *sucA, sucB*, and *lpdA*, which converts akg to succoa while generating NADH (Fig. 8A-B). Gene expression of *sucAB* in all eSdhCB strains was upregulated compared to uSdhCB except for eSdhCB01, which was significantly depressed compared to all other strains. The drop in gene expression of eSdhCB01 was most likely due to an intergenic mutation (Fig. 8C, Table S4). In addition, flux appeared to be reduced in all strains due to mutations that affected substrate binding and complex formation (Fig. 8C). A DEL mutation in *sucA* in eSdhCB03 occurred at residues that affect substrate binding, while SNP mutations in multiple strains occurred in regions that could affect *sucA* homomer formation (Fig. 8D). A DEL mutation in *sucB* in eSdhCB01 cleaved residues 405 to 273, which are located in the active site (Fig. 8D). In total, these mutations appeared to confer a fitness advantage in all eSdhCB strains by modulating the flux through AKGDH, which ultimately reduced succ byproduct secretion.

**Fig. 8.**
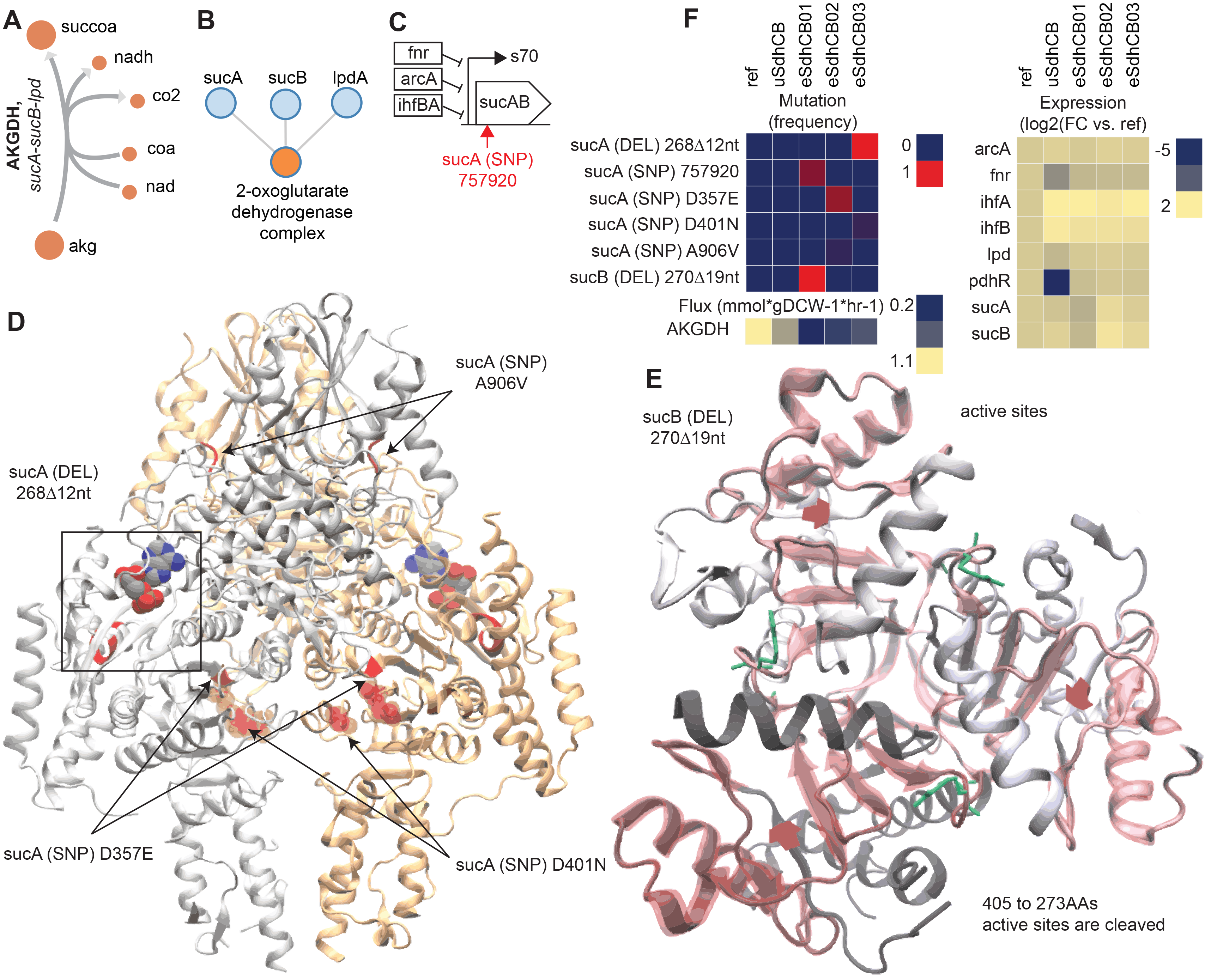
Mutations that affect 2-oxoglutarate Dehydrogenase (AKGDH) enzymatic activity and expression in eSdhCB strains. A) Network diagram of the AKGDH reaction. B) Protein-protein binding diagram of the 2-oxoglutarate Dehydrogenase complex formation. C) Operon diagram of the *sucAB* operon. The position of the intergenic mutation is highlighted in red. Gene expression of *sucAB* in eSdhCB01 is significantly depressed compared to all other strains, most likely due to the intergenic mutation (Table S4). In contrast, *sucAB* is upregulated in eSdhCB02/03 compared to uSdhCB (Table S4). D) Crystal structure of SucA. Mutations are highlighted in Red. A substrate analog is shown. The deletion (DEL) mutation in eSdhCB03 occurred at residues that affect substrate binding. The sucA D357E, A906V, and D40N SNPs all occur near homodimer interface regions. E) Crystal structure of SucB. Mutations are highlighted in red. Cleaved regions are outlined in red. Substrate binding residues are highlighted in green. The DEL mutation cleaved residues 405 to 273, which are all in the active site. F) Mutation frequency, gene expression levels, and flux of components involved in the AKGDH reaction. Note the reduced flux through AKGDH in all eSdhCB strains.

### Adaptive evolution re-wired the flux map of eGnd and eSdhCB strains despite an insignificant change in growth rate

While insignificant changes in growth rate were found in most eGnd and eSdhCB strains, massive changes in metabolic flux were found (Table S6). These shifts in flux were a result of mutations that were selected for during evolution (Figs. 3-8, Table S8) as described above. For example, while flux was initially diverted through the ED pathway in uGnd, all eGnd strains completely inactivated oxPPP and ED flux in exchange for increased flux through upper glycolysis and the nonOxPPP (Fig. 2A-B). This shift also entailed another reversion of flux through the transketolases and transaldolase reactions (Fig. 2A-B). The inactivation of the oxPPP was made possible by increased flux through ICDHyr and mutations in THD2pp (Fig. 4) that made up for the lack of NADPH production in the oxPPP. Major flux shifts were also found in all eSdhCB strains. Most notably, TCA cycle flux was significantly reduced in all eSdhCB strains (Fig. 2C-D). This shift was made possible by mutations affecting nitrogen metabolism (Figs. 5-6), sulfur metabolism (Fig. 7), and AKGDH in the TCA cycle (Fig. 8) that appeared to coordinate the amount of flux out of the TCA cycle with generation of precursor metabolites derived from the TCA cycle.

## Conclusion

While disruption of the GND and SUCDi reactions in the oxPPP and TCA cycles, respectively, resulted in minimal changes to growth rate, massive changes in metabolite levels and metabolic flux were found. Interestingly, many similarities in gene expression states between both strains despite the large difference in network location of the perturbations were found. Commonalities in gene expression profiles could be traced back to known metabolite regulators that were similarly perturbed in both strains. Minimal changes in growth rate were found after ALE, but mutations that lead to major divergence in flux and gene expression states from the unevolved strains were found. In the *gnd* evolutions, mutations were found that compensated for the reduced ability to generate NADPH through the oxPPP; in the *sdhCB* evolutions, mutations were found that reduced TCA cycle flux while balancing the regulation of nitrogen and sulfur assimilation. The divergence of regulatory states after evolution demonstrates that while metabolic and regulatory networks are robust to perturbation, the initial adjustments are often not optimal even when minimal changes in fitness occured. It is only after adaptation that the optimal regulatory and flux states were revealed.

## Contributions

D.M. designed the experiments; generated the strains; conducted all aspects of the metabolomics, fluxomics, phenomics, transcriptomics, and genomics experiments; performed all multi-omics statistical, graph, and modeling analyses; and wrote the manuscript. T.E.S. ran the ALE experiments. E.B. assisted with structural analysis. R.S. processed the DNA and RNA samples. S.X. assisted with metabolomics and fluxomics data collection, sample processing, and peak integration. Y.H. assisted with fluxomics data collection and sample processing. A.M.F designed and supervised the evolution experiments, and contributed to the data analysis and the manuscript. B.O.P conceived and outlined the study, supervised the data analysis, and co-wrote the manuscript.

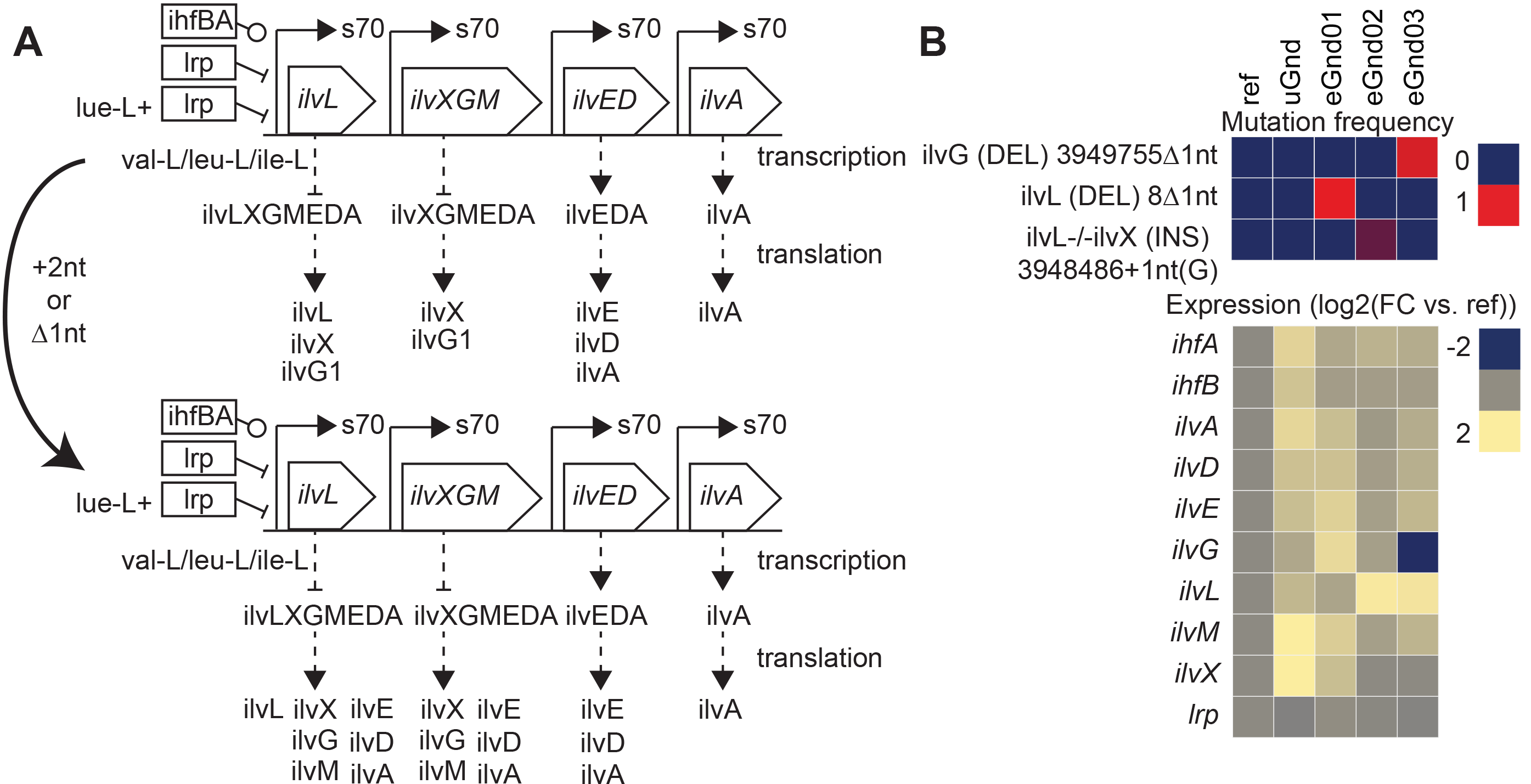

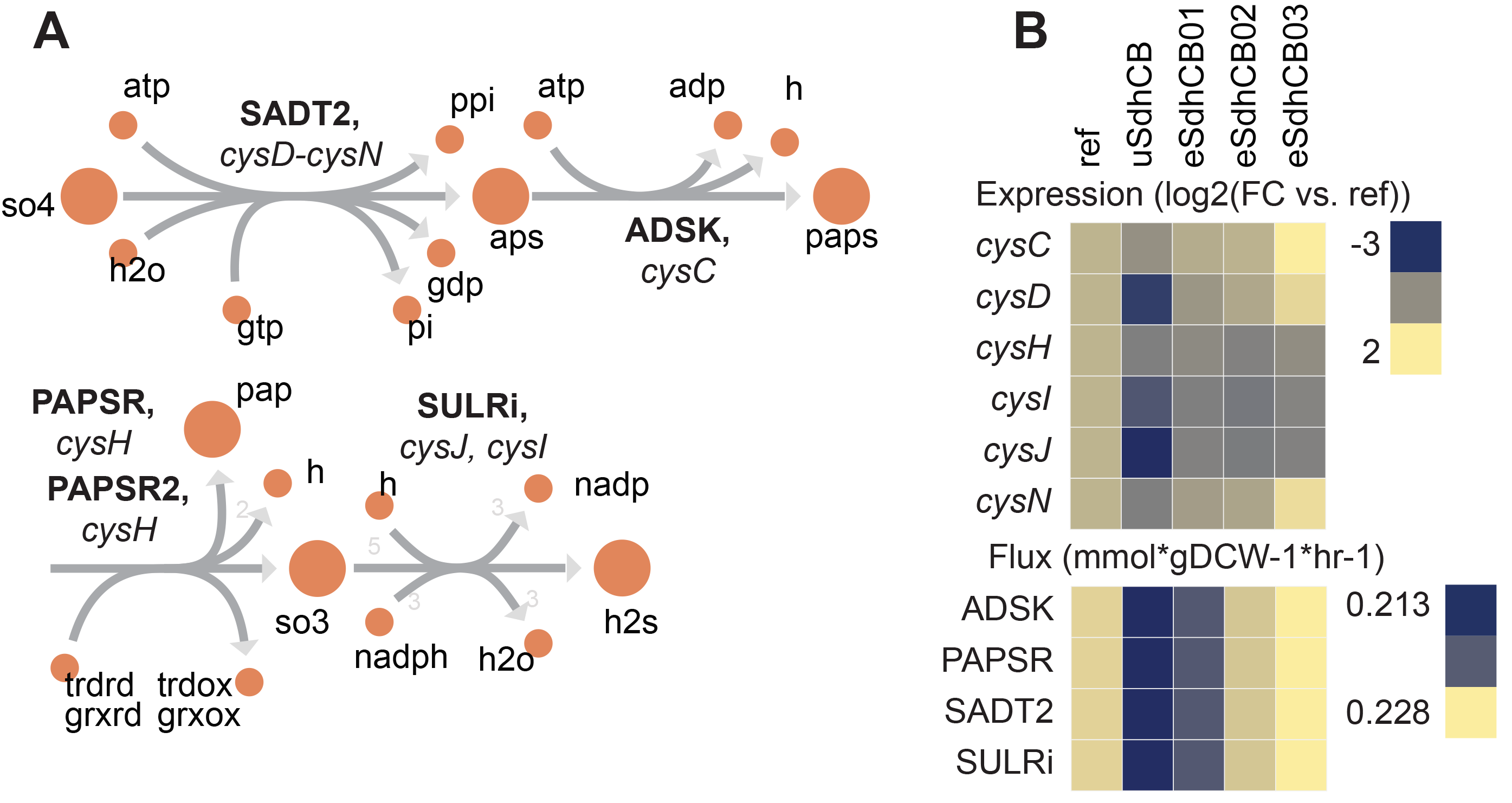

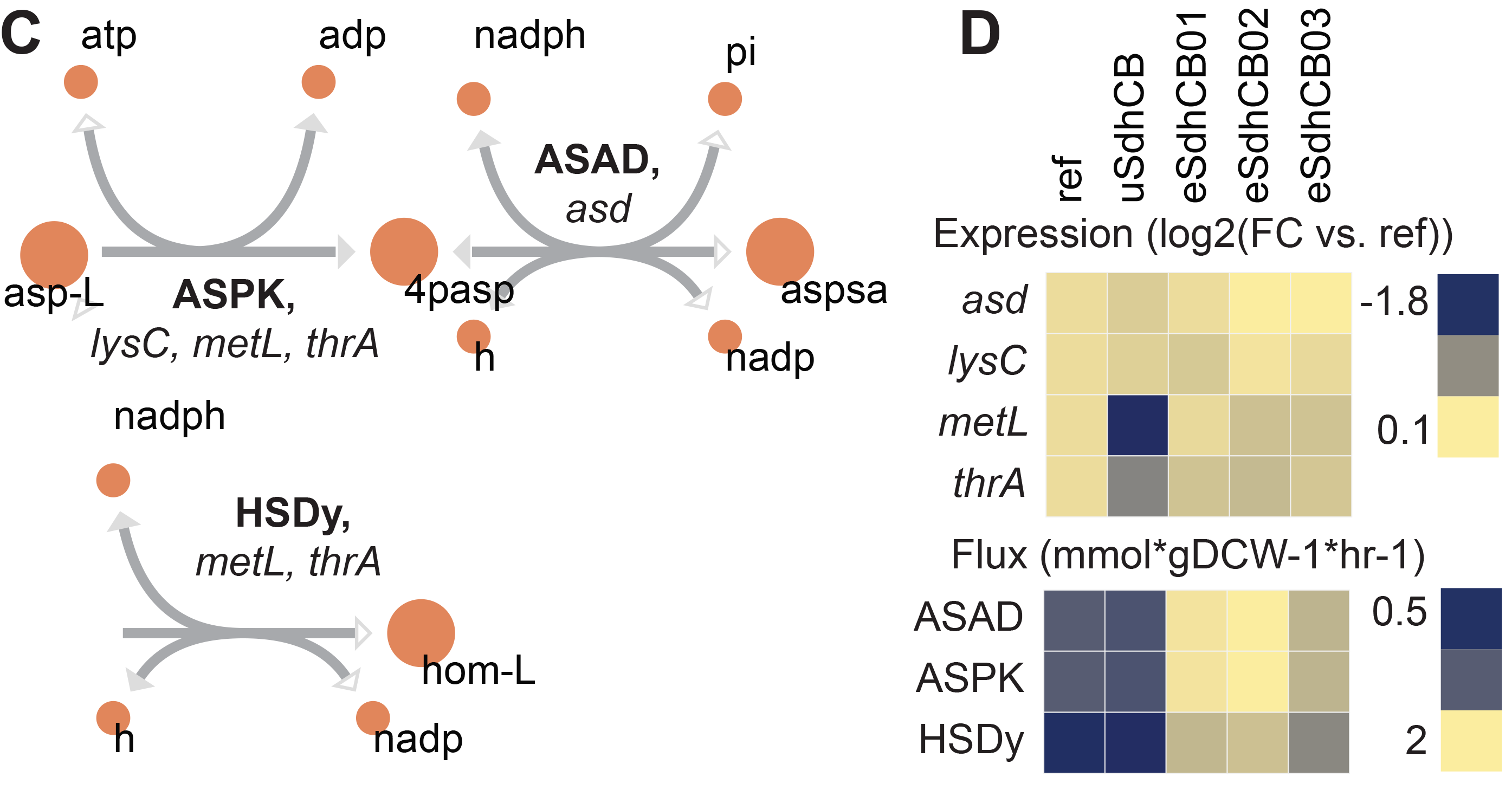

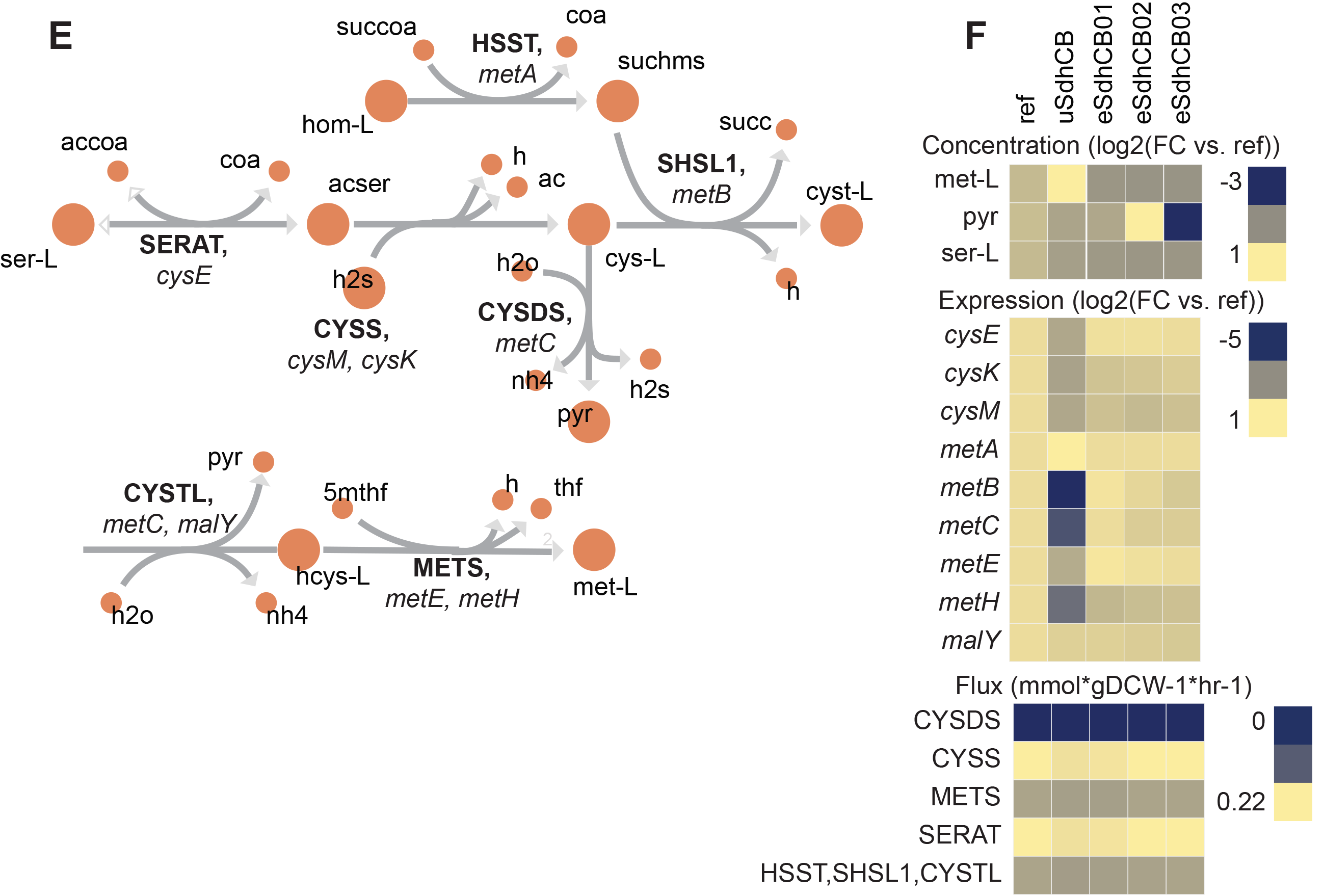

## Acknowledgements

We thank José Utrilla for helpful discussion and guidance when implementing the knockouts in the pre-evolved strain. We thank Jamey Young for helpful discussions throughout the MFA analysis. This work was funded by the Novo Nordisk Foundation Grant Number NNF10CC1016517.

